# Geometry-Optimized Inkjet-Printed Organic Electrochemical Transistors with High Transconductance and Sub-Millisecond Response for Biosensing and Neural Recording

**DOI:** 10.1101/2025.11.01.686019

**Authors:** Fadi Khoury, Zeina Habli, Jad Daorah, Yuchen Xu, Makram Obeid, Samir Alam, Gert Cauwenberghs, Massoud Khraiche

## Abstract

Organic Electrochemical Transistors (OECTs) are witnessing rapid growth in biomedical applications and are increasingly becoming an integral part of bio-electronic interfaces. High-performing OECTs are typically fabricated using multistep photolithography and conventional spin-coating and lift-off processes, and while printing techniques have emerged as promising alternatives, they still face challenges in achieving comparable resolutions, reproducibility and performance metrics. Several groups have demonstrated printed OECTs using PEDOT:PSS as the channel material, highlighting the promise of additive manufacturing for scalable bioelectronics. In this work, we build upon these advances and develop an optimized inkjet-printed OECT platform that achieves transconductance values up to 15 mS and sub-millisecond response times as low as 0.31 ms. Our approach systematically optimizes OECT geometrical parameters—channel width, length, and thickness—through precise patterning and oxygen plasma surface modification to overcome longstanding limitations in inkjet printing resolution and reproducibility. The resulting devices exhibit outstanding electrical stability, high amplification, and fast dynamic response. Using a configuration optimized for biosensing, we demonstrate the detection of the heart failure biomarker NT-proBNP within a clinically relevant range of 10–400 pg/mL, with a sensitivity of 0.038% ΔI_DS_/pg/mL. In a separate configuration on a flexible substrate tailored for in vivo biopotential recording, we showcase the devices’ capabilities by effectively capturing epileptic seizure progression in a rat model with high signal fidelity. This work demonstrates how careful process and geometry optimization can close the performance gap between printed and conventionally fabricated OECTs, enabling scalable, reproducible, and substrate-flexible bioelectronic platforms.

## Introduction

The field of bioelectronics, valued at 7.77 billion dollars in 2022 and projected to reach a market size of 13.86 billion dollars by 2030, is undergoing rapid growth fueled by the growing demand for advanced medical technologies for personalized and effective treatments^1^. This surge is supported by the advancements in bio/electro-chemistry, surface chemistry, electronic devices, and materials science, leading to the development of new and innovative bioelectronic devices. Central to this technological shift in biosensor research are Electrolyte-gated transistors (EGTs), which are currently used in a many of biosensing applications including infectious diseases, biomolecule sensing, and electrophysiological monitoring, showing high sensitivity and low cost ^2–6^. Organic Electrochemical Transistors (OECTs), invented around three decades ago^7^, are types of EGTs^8^, and have emerged as pivotal tools in biosensing given their impressive compatibility with the biological milieu. Organic mixed ionic-electronic conductors (OMIECs) are used as active channels and allow direct channel-electrolyte contact, employing mixed ionic/electronic charge transport for direct transduction of biological activity to measurable electrical signals in the form of drain current flow^9,10^. OECTs, particularly those based on the semi-conducting polymer poly(3,4-ethylenedioxythiophene): polystyrene sulfonate (PEDOT: PSS), stand out given their ability for bulk channel de/doping, which allows for the modulation of charge carrier density throughout their bulk. This results in superior capacitive coupling between the electrolyte and the channel, characterized by the increase in capacitance with increased channel thickness (i.e. volumetric capacitance). Additionally, PEDOT:PSS sets itself apart from other OMIECs by having a relatively high hole mobility^11–13^. These key characteristic features enable PEDOT:PSS-based OECTs to achieve milli-siemens order transconductances (g_m_), rendering them effective for low biomolecule concentration detection and low voltage signal amplification^14,15^. While most of the research on OECTs has been focused on their operation, application, and engineering of next-generation channel materials^16–18^, some effort has been made in the exploration of reliable and accessible fabrication methods to expand the use of OECTs without being constricted to specialized clean room facilities. The latter requires the use of harsh chemicals for photolithography and wet etching, in addition to high-temperature processes for metal deposition, limiting the choice of substrate materials and might also pose risks to heat-sensitive components, limiting integration^19^. In addition, these procedures raise concerns on expenses/cost, waste materials, and complexity. Although methods such as inkjet printing, screen printing, aerosol jet printing, and various combinations of these printed techniques have been previously used to fabricate OECTs, they fail to achieve dimensions in the low micrometer scale for lengths and the low nanometer scale for thicknesses, important for unlocking performance metrics suitable for a wide range of applications in biomolecule detection and in electrophysiological monitoring ^20–40^. In this work, we address these limitations by optimizing the inkjet printing process via surface modification to precisely control the thickness, width, and length of PEDOT: PSS channels. We also systematically examine how these geometrical parameters influence electrical behavior and demonstrate device versatility in two critical biomedical contexts: (1) biosensing of the NT-proBNP heart failure biomarker, and (2) in vivo electrocorticography (ECoG) recording in a rodent seizure model. Our findings provide a fabrication framework for producing highly stable, reproducible, and clinically relevant OECT-based bioelectronic devices, with potential applications spanning point-of-care testing and flexible neurointerfaces.

## Methodology

### Device Fabrication

PEDOT:PSS OECTs were fabricated on clear microscopic glass slides using the Dimatix Materials Printer (DMP-2850, Fujifilm, Dimatix, USA) fitted with a 2.4 pL piezo-driven 12-nozzle printhead (Samba, PDS00142). Briefly, the glass slides were cleaned with ethanol and thoroughly rinsed with Milli-Q water and placed at 250°C in a drying oven for 30 min before use. Next, the OECTs were fabricated with three main steps: (1) deposition of gold nanoparticle conductive ink (JG-125, Novacentrix, USA), 1 layer at 20 µm Drop Spacing (DS), to form the source (S) - drain (D) leads and the contact pads, followed by sintering at 180°C for 35 minutes in a drying oven. (2) insulation of the gold interconnects to realize the active channel with polyimide ink (UT Dots-PI-IJ) at 4 layers, 15 µm DS, and curing at 180°C for 1 hour. To avoid variability in the electrical characteristics of the fabricated devices due to channel–electrode overlap, the overlap was maintained at 30 ± 5 μm for all devices. (3) printing the active channel with inkjet-printable PEDOT:PSS aqueous conductive ink (0.8% in H2O, 739316, Sigma Aldrich, USA) in 1 to 3 layers, at either 15 or 25 µm DS to connect the S&D, which was cured at 150°C for 20 minutes. At least three devices were printed for every configuration for accurate representations of device performances across fabrication iterations, all of which shared the same interconnect dimensions to eliminate series resistance variability. For devices fabricated with complementary oxygen plasma treatment, dry oxygen-etching treatment was performed prior to PEDOT:PSS deposition, for a duration of 3 or 10 s, using a plasma etcher (PE-25, PlasmaEtch) at 500 mTorr vacuum set point, 150 W RF power, and 15 cc/min oxygen flow rate.

### Electrical Characterization

Electrical recordings were performed using source measuring units SMUs (PXIe-4138, National Instruments, USA). All electrical characterizations were carried out in 1X Phosphate Buffer Saline (PBS, P4417, Sigma) using an Ag/AgCl wire as the gate electrode. To acquire the Current-Voltage (IV) scans for all intended configurations, the source was grounded, the drain-source voltage was swept from 0 to −0.6V (100 points), while the gate-source voltage was stepped (10 steps) from 0 to +0.6 V. For transfer characteristic scans, the drain-source voltage was fixed at −0.4 V, but was only set at −0.6 V for repetitive scans to assess the stability of the OECTs in the saturation window (from 0.1 V_GS_ to 0.45 V_GS_). The gate-source voltage range was not fixed for all devices and depended on the threshold voltage to ensure proper saturation in both the fully conductive and fully insulating regimes. To extract the response time of the devices, a square wave was applied to the gate terminal with a 50% duty cycle and a desired frequency of 1 Hz or 20 Hz, depending on the ability of the device’s switching speed for good quality fits. The limits of the wave depended on the limits of the transfer characteristics of the corresponding device to ensure maximum and minimum current flow. The drain current output was recorded at a 2500 Hz sampling frequency, and the transition from the On-state (maximally conducting) to the Off-State (minimally conducting) was fitted to an exponentially decaying function with a time constant representing the response time ( ), following the methods reported in the work of Ohayon et.al. ^15^. The cut-off frequency is then deduced by calculating the inverse of . In the artificial neuron spike measurements, the 60MEA2100-SG (population spike signal) from Multichannel Systems was connected to gate terminal, offsetted by V_GS_ through a series connection with the an SMU, and was grounded to the source. The SNR (in dB) was calculated by taking the ratio of the maximum amplitude peak and the standard deviation of the baseline noise where no response is observed.

### PEDOT:PSS Cross-Sectional Height Measurements

PEDOT:PSS was either printed on cleaned glass on 10s oxygen plasma-treated glass at 15 µm DS, on cleaned glass at 25 µm DS, or on 2s plasma-treated glass at 25µm DS resembling OECT channels of different sizes and thicknesses. Digital holographic microscope (DHM®, LyncéeTec, Switzerland) was used to measure the vertical and horizontal cross-sectional dimensions of PEDOT:PSS prints using holographic interferometry technology to produce live holographic 3D reconstructions of the specimen for cross-sectional profiling in an optical non-contact point.

### Form and Function: Influence of Geometry on OECT Characteristics

dimensions of OECTs are critically influenced by their intended application. The latter plays a decisive role in shaping the geometric attributes of the semi-conducting channel, directly affecting the transconductance (g_m_) of a depletion mode OECT (as is the case for hole-conducting PEDOT:PSS) following:

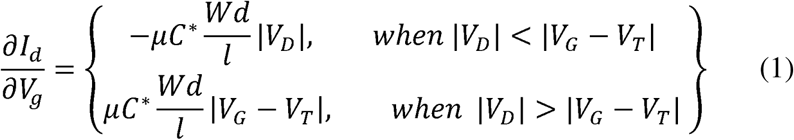

Where: *I_d_* is drain current; *V_G_* is gate voltage; μ is the hole mobility; *C*\* is the volumetric capacitance; *W* is channel width; *d* is channel thickness; *l* is channel length; *V_D_* is the drain voltage; *V_T_* is threshold voltage12. In addition, the geometry contributes directly to modulating the response time, following

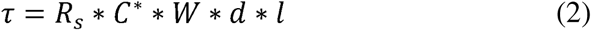

Where: *τ* is the response time; *R_S_* is the electrolyte resistance in turn proportional t 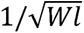.

Equation (1) and (2) highlight the intricacy between the device form and its intended function, such that the design of the OECTs must be tuned to fulfill application-specific needs ^11,18^.

### Gate Functionalization and NT-proBNP Detection

To functionalize the gold with antibodies the following steps were taken. The gold electrode is immersed in SH-PEG-Biotin (100 µg/ml) for 2 hours, the thiol groups have high affinity for gold, realizing a strong covalent bond. Next, streptavidin (100 µg/ml) is added to the electrode surface and is incubated for one hour. Following the streptavidin incubation, the electrode is washed again with PBS to remove any unbound streptavidin. Next, the biotinylated antibodies (100 µg/ml) are added to the electrode surface for a duration of one hour. Finally, to minimize non-specific binding and improve the specificity of the functionalized surface, 100 µg/ml of Bovine Serum Albumin (BSA) is introduced to the electrode surface. After every incubation period, the electrode is thoroughly rinsed with phosphate-buffered saline (PBS) to remove any unbound molecules.

Calibration curves were generated using three devices, with incremental additions of NT-proBNP concentrations (10–1000 pg/mL). Transfer curves were recorded 5 minutes after introducing each protein sample. Similarly for controls, equivalent volumes of buffer-only aliquots were added, and transfer curves were measured after 5 minutes to assess the natural drift of the current (I_DS_) over time. A shift in I_DS_ exceeding the baseline drift was considered a response to protein concentration. The shift, ΔI_DS_, was calculated as ΔI_DS_ = I_DS, [protein]_ − I_DS, [0]_, where I_DS_ is measured at the gate-source voltage corresponding to the maximum transconductance.

### In-Vivo Experiments

Animal Experiments were approved by the Institutional Animal Care and Use Committee (IACUC) at the American University of Beirut (Form # 001/05). Adult male Sprague-Dawley rats, weighing 300-350g (n=3 animals) were used for electrocorticography recordings. Anesthesia was administered intraperitoneally with ketamine (80 mg/kg) and xylazine (20 mg/kg). The craniotomy is performed over the somatosensory cortex and is about 4 x 4 mm on the left side of the brain. A surgical drill is then used to drill off the bone, after which the dura is removed, exposing the brain surface. To induce seizure activity, a 1 μL Hamilton syringe is fixed to a stereotaxic holder, 0.6 μL of 1mg/ml kainic acid (KA) in saline is then injected into the amygdala (−2.8 mm Anteroposterior (AP), 5 mm Mediolateral (ML), and 8.8 mm Dorsoventral (DV) referenced to the bregma). The OECT flexible device is then placed over the cortex and connected to the ME2100-system (Multichannel Systems). More details can be found in our previous publications ^41^.

## Results

In this work, we explore the feasibility of OECT devices fabricated with Inkjet printing for multipurpose applications as demonstrated in figure 1a. Three different channel geometric parameters of the OECT channels - width (W), thickness (d), and length (l) were investigated to elucidate their impact on OECT performance (fig. 1b) and set some strategic key points for fabricating inkjet-printed OECTs as bio-electronic interfaces. The modifications included either adjusting the dimensions of the source and drain, fine-tuning the thickness of the active channel, or narrowing the gap between the source and drain. Subsequent electrical measurements were conducted to assess the performance and implication of these variations.

**Figure 1.**
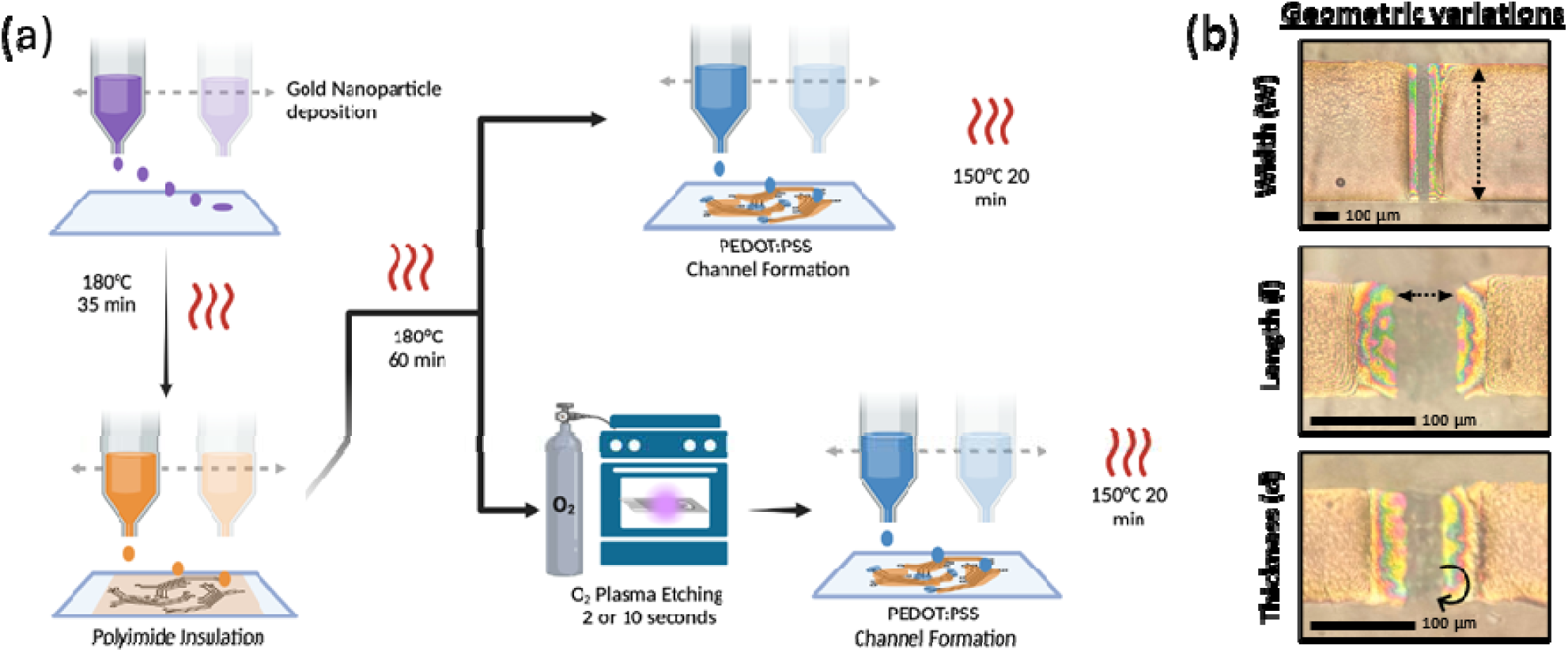
(a) Inkjet printed OECT fabrication flow with complementary oxygen plasma treatment allowing for the precise control of the three OECT geometrical parameters; **(b)** Width (W), Length (l), and Thickness (d).

### Width variation

In this section, for a fixed channel length (l = 40 µm) and channel thickness (d = 2 layers equivalent to 400 nm), three different channel widths were configured: 100 µm, 300 µm, and 500 µm. These OECT configurations will be referred to as W100, W300, and W500, respectively. The DHM analysis of the designed widths revealed accuracies of 95.2%, 91.5%, and 97.8%, respectively, with minimal standard deviations (fig.s2) confirming high width accuracy. Analysis of the electrical characteristics of the configured OECTs reveals that as the channel width increases, there is a corresponding increase in the current levels flowing through the channel, particularly at low gate voltages (fig.2a-2c). Additionally, transfer characteristics indicate higher on/off ratios and a noticeable positive shift of the active region for the larger channels, resulting in increased threshold voltage (fig.2d) in accordance with previously reported findings ^42^. Notably, the maximum g_m_ reached was 12.5±0.4 mS (25 mS/mm) for W500, compared to 9.9±0.9 mS (33 mS/mm) for W300 and 4.3±0.12 mS (43 mS/mm) for W100. Similarly, the location of maximum g_m_ increases with width, falling at 0 V_GS_, 0.12 V_GS_, and 0.26 V_GS_ for W100, W300, and W500, respectively (fig.2e). Conversely, the response time, and consequently cut-off frequency, decreases with increasing width. This is evident when recording the output responses of the devices under a square wave applied at the gate, showing that the larger channels failed to accurately reconstruct the full wave at higher frequencies (fig.S1a). Fitting the device’s responses to the 1 Hz square wave yielded response times of 1.5±0.2 for W100, 3.9±0.3 for W300, and 4.8±0.7 ms for W500, resulting in cut-off frequencies of 107 Hz, 41 Hz, and 34 Hz, respectively (fig.S1b-1d). For assessing stability, the W500 device was subjected to 100 consecutive scans in the transfer characteristic’s saturation regime at −0.6 V_DS_; the latter will increase the g_m_ from 12.5 mS to 15 mS compared to the −0.4 V_DS_ setting. The W500 device displayed high stability and overlapped across scan cycles(fig.2f).

The standard deviation of the current as a function of gate voltage reveals the device’s most stable point which coincides with the maximum g_m_ location, indicating an error margin of ∼ 40 µA compared to a maximum of ∼90 µA at either end of the transfer characteristic. Noticeably, g_m_ increased slightly with scanning repetitions (fig.2-g) from 15 mS to 15.7 mS by the 100^th^ scan. In fact, beyond 50 scans, all three studied parameters (g_m_, the location of g_m,max_ (V_GS, gm max_), and the current levels at g_m,max_ (I_DS,gm max_)), began showing slight signal drifts, indicating some minor instabilities (fig.2h). These findings underscore the critical role of channel width in optimizing OECT performance, highlighting potential for tailored designs to significantly enhance the device’s signal fidelity and stability for bio-electronic applications.

**Fig. 2.**
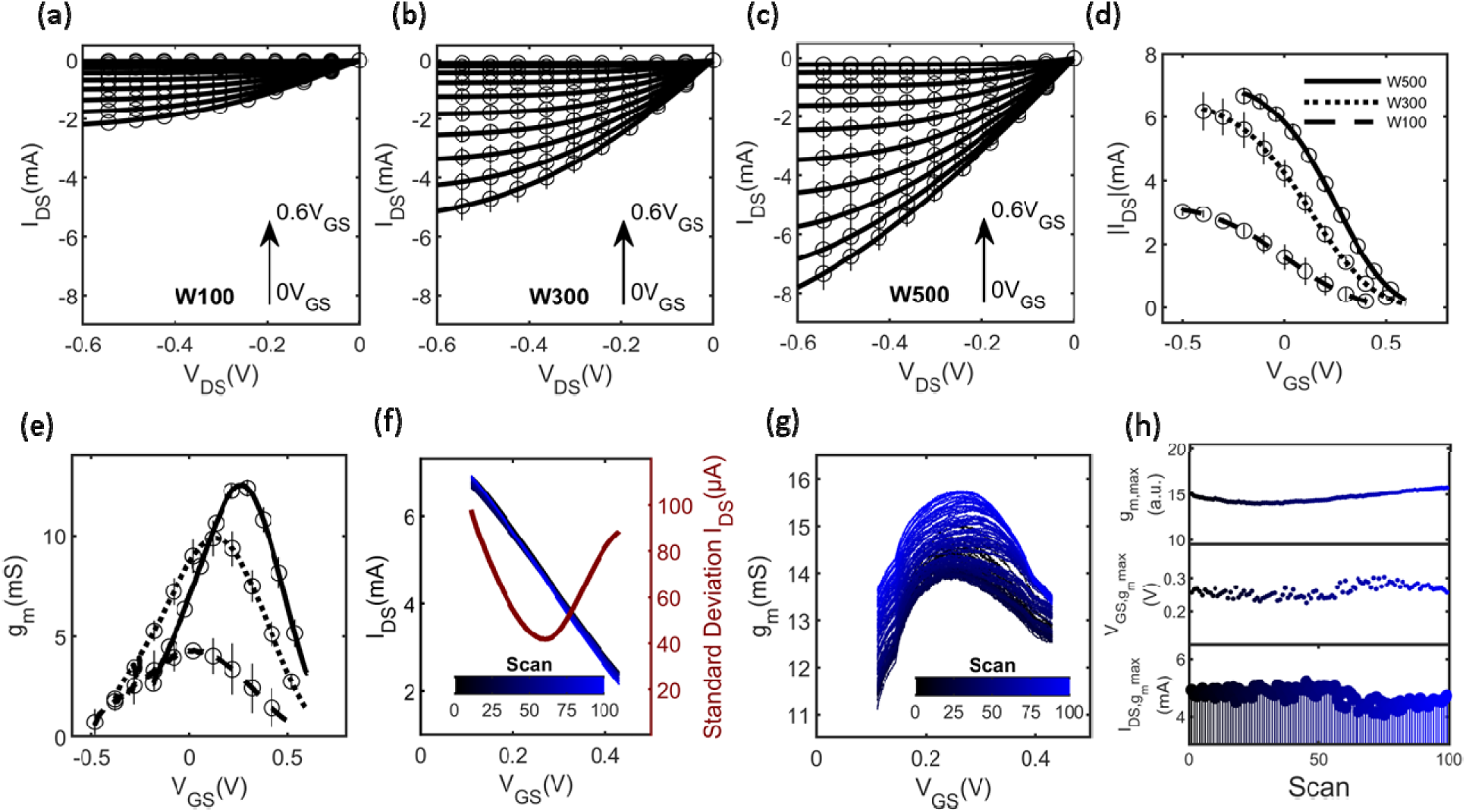
IV curves of **(a)** W100, **(b)** W300, **(c)** W500 with VDS from 0V to −0.6V and VGS from 0V to 0.6V. **(d)** transfer characteristic and **(e)** gm curves of W100 (dashed), W300 (dotted), and W500 (solid line) with VGS from −0.5V to 0.6V and −0.4VDS. **(f)** repetitive transfer characteristic scans (100) in the saturation window (0.1V_DS_ to 0.45V_DS_) of W500 at −0.6V_DS_ and the corresponding standard deviation in red. **(g)** associated g_m_ curves of the repetitive scans. **(h)** variation of maximum g_m_ (top) V_GS, gm max_ (middle) and I_DS, gm max_ (bottom) across electrical scans.

**Fig. 3.**
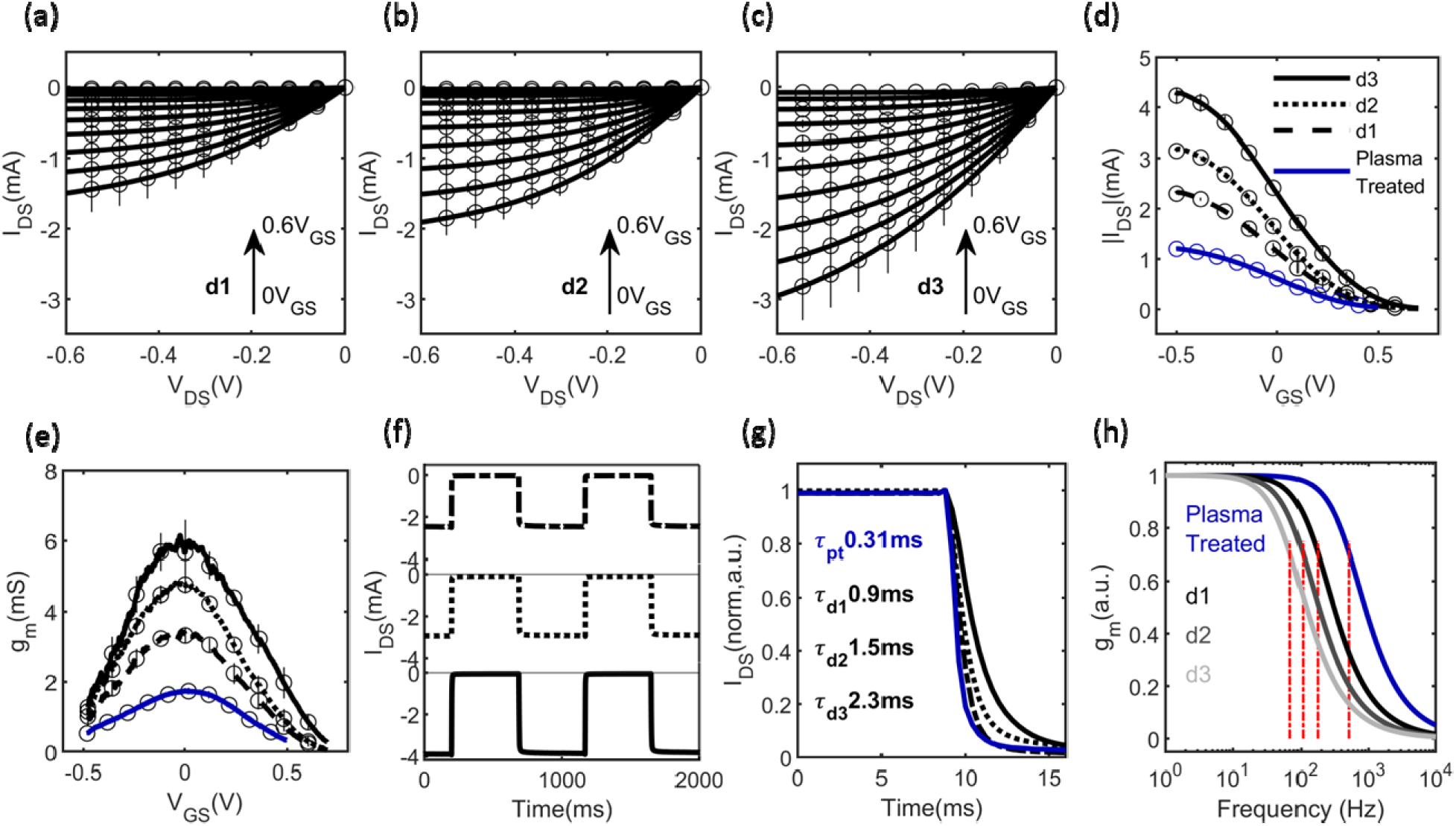
IV curves of **(a)** d1, **(b)** d2, **(c)** d3 with VDS from 0V to −0.6V and VGS from 0V to 0.6V. **(d)** transfer characteristic and **(e)** gm curves of d1 (dashed), d2(dotted), d3 (solid), and plasma treated (solid blue line) with VGS from −0.5V to 0.6V and −0.4VDS. **(f)** current response of d1 (top), d2 (middle), and d3 (bottom) under a 1 Hz square wave applied at the gate with limits ensuring upper and lower boundary saturation. **(g)** zoomed in normalized response of d1 (dashed), d2(dotted), d3 (solid), and plasma treated (solid blue line) from which response time (τ) is extracted by fitting the data to a decaying exponential curve. **(h)** modeled normalized g_m_ amplitude response to frequency of d1 (black), d2(grey), d3 (light grey), and plasma treated (blue).

### Thickness variation and oxygen plasma treatment

Next, three thicknesses of PEDOT:PSS were investigated for OECTs: d1 with one layer of 200 nm thickness (measured at 212±18 nm), d2 with two layers of 400 nm thickness (measured at 417±38 nm), and d3 with three layers of 600 nm thickness (measured at 609±65 nm), while maintaining a fixed channel width (W = 100 µm), and length (l = 40µm). Analysis of the IV output curves demonstrate that higher current values flow through PEDOT:PSS with increased thickness (fig.3a-3c), alongside higher on/off ratios as indicated by the transfer characteristics (fig.3d). Interestingly, while the maximum g_m_ values scale up with increased PEDOT:PSS thickness - from 3.4±0.2 mS (34 mS/mm) for d1 to 4.8±0.2 mS (48 mS/mm) for d2 to 6.1±0.9 mS (61 mS/mm) for d3 - there was no observed lateral shift in maximum g_m_, contrary to width variation results (fig.3e). Notably, devices with thinner channels exhibited faster response times, scoring 0.9±0.17 ms, 1.5±0.2 ms, and 2.3±0.5 ms (fig.3f and fig.3g), and higher mean cut-off frequencies: 177 Hz, 107 Hz, and 70Hz for d1, d2, and d3 respectively (fig.3h).

Next, we tune our fabrication to push the limits of inkjet printing in thickness. We achieve this by modifying the substrate surface with oxygen plasma before depositing PEDOT:PSS, rendering the surface more hydrophilic, improving film uniformity. First, three one-layer PEDOT:PSS squares of sizes 120×120 µm (8×8 pixels), 300×300 µm (20×20 pixels), and 500×300 µm (33×33 pixels) were printed on a non-treated glass substrate at 15 µm DS.

All the prints had a standard thickness of around 212±18 nm extracted by measuring the cross-sectional profiles at the horizontal midline using the DHM. However, plasma treatment of glass for 10s prior to printing reduced the necessary volume of PEDOT:PSS by 57±7% to realize the same print sizes with a thinner thickness of 129±31 nm, a 0.4-fold decrease compared to non-oxygen plasma treated glass (fig.4b). If not for oxygen plasma treatment, reducing the volume of deposited ink would result in diminished coverage, exposing the metal contacts to the electrolyte. Nonetheless, meticulous management of surface wetting with oxygen plasma treatment ensures the preservation of the print’s spatial footprint while exclusively reducing its thickness. Another parameter to be tested was raising the DS to 25 µm and reducing oxygen plasma treatment time to 2s; this allowed for printing uniform PEDOT:PSS squares with full coverage of the desired area using only one layer and achieving a significantly reduced thickness of 45±15 nm, an attribute not achievable by simply increasing the DS only where several gaps in the prints were reported even with 2 consecutive layers as seen in figure 4c. However, OECTs fabricated with this thin film thickness (45±15 nm), were unstable under applied bias, due to poor adhesion of PEDOT:PSS to the underlying gold and glass substrate, especially when biased. Thus, to guarantee stability, reproducibility and high yield, we opted to work with devices having 129 nm PEDOT thickness.

**Fig. 4.**
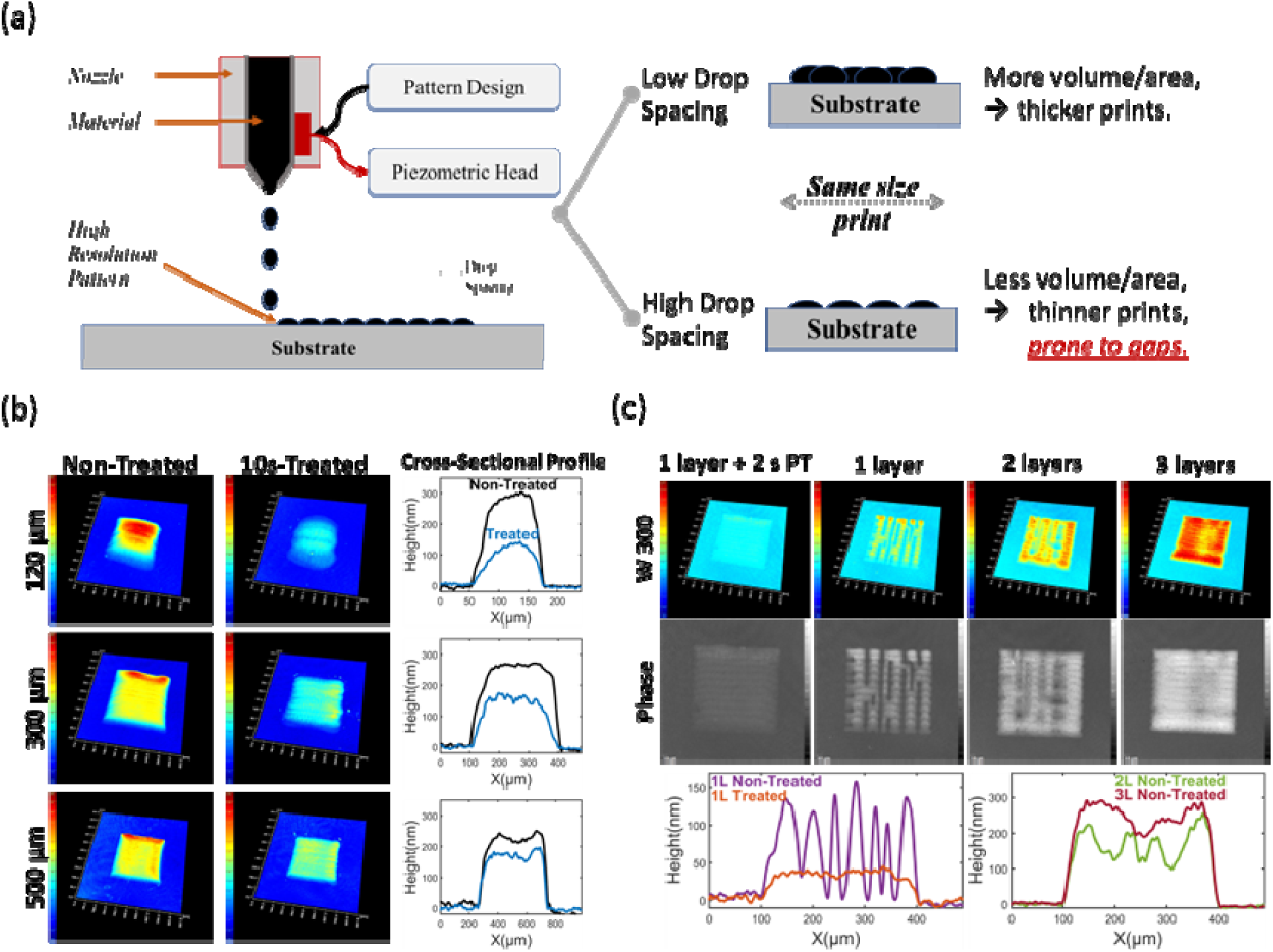
(a) Schematic showing the effect of drop spacing in inkjet microfabrication on print thickness. (b) Comparing the DHM images and cross-sectional profiles of PEPOT:PSS 1-layer squares deposited on non-oxygen plasma treated glass and 10-second oxygen plasma treated (PT) glass at DS15µm and (c) Comparing the DHM and Phase images and cross-sectional profiles of PEPOT:PSS squares deposited on non-oxygen plasma treated glass (1 or 1L, 2 or 2L, and 3 or 3L layers) and 2-second PT glass (1 layer) at DS25µm.

**Fig. 5.**
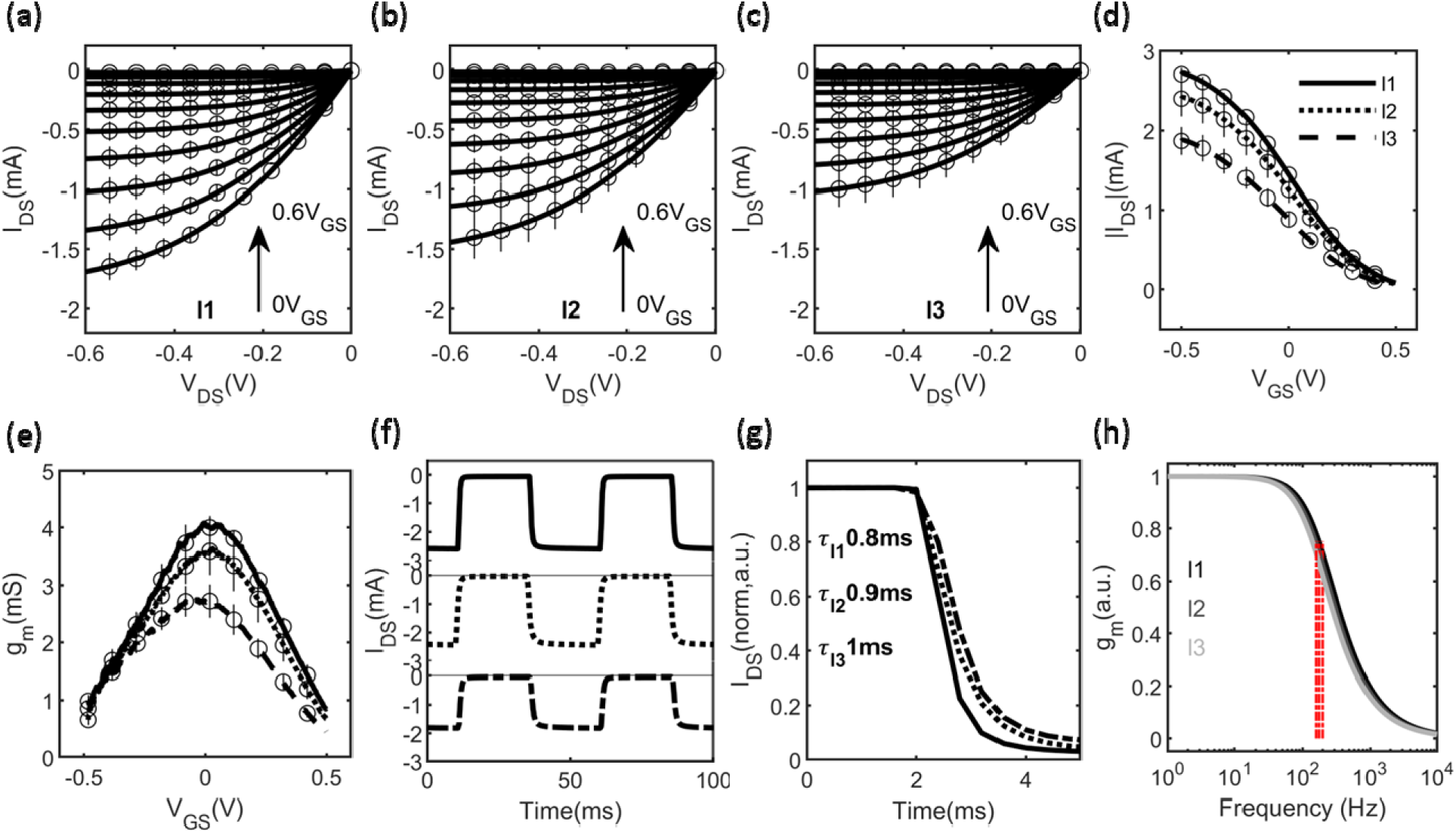
IV curves of **(a)** l1, **(b)** l2, **(c)** l3 with V_DS_ from 0V to −0.6V and V_GS_ from 0V to 0.6V. **(d)** transfer characteristic and **(e)** g_m_ curves of l1 (solid), d2(dotted), and d3 (dashed) with V_GS_ from −0.5V to 0.5V and −0.4V_DS_. **(f)** current response of l1 (top), l2 (middle), and l3 (bottom) under a 20 Hz square wave applied at the gate with limits ensuring upper and lower boundary saturation. **(g)** zoomed in normalized response of l1 (solid), l2(dotted), and l3 (dashed) from which response time (τ) is extracted by fitting the data to a decaying exponential curve. **(h)** modeled normalized g_m_ amplitude response to frequency of l1 (black), l2(grey), and l3 (light grey).

A challenge evident in the phase images and the corresponding thickness profiles indicates that a high DS before oxygen plasma treatment leads to incomplete non-uniform layer formations. These gaps can only be filled by increasing the number of printed layers to two or even three, thus increasing the total thickness of the channel, and counteracting the objective of increasing the response time as seen in the cross-sectional profile scans in figure 4c. To verify the reliability of the proposed method, modified OECT devices (n=3) were realized with a channel thickness of 129±31 nm (using the earlier oxygen plasma treatment method of DS 15 µm coupled with 10 s of oxygen plasma treatment prior to printing), a width of 100 µm, and a length of 40 µm were tested. Compared to d1, IV curves show a decrease in current flow at every V_GS_ step along the sweep of V_DS_ (fig.S3a), and a decrease of the on/off ration (fig.3d, blue line), alongside a reduction in g_m_ to 1.7±0.1mS equivalent to 17 mS/mm (fig.3e) as expected by a thinner channel. However, oxygen plasma treatment and DS modifications led to marked decrease in response time (fig.3g and fig.S3b), and cut-off frequency (fig.3h) to 0.31±0.01 ms and 514 Hz, respectively, showcasing the nuanced impact of PEDOT:PSS layer thickness on device performance.

### Length Variation

Finally, to examine the effect of channel length and its limits in Inkjet microfab, three OECT configurations were fabricated maintaining a constant width (W = 100 µm) and channel thickness (d = 200 nm), but with varying lengths: l= 20 µm (l1), 40 µm (l2), and 60 µm (l3). The channel lengths were verified using DHM, measuring 19±1 µm, 38±2 µm, and 56±2 µm respectively (fig.s2). The IV curve analysis shows that larger currents flow in the channel (fig.5a-5c) with decreasing lengths, contrary to the trends observed with increasing both width and thickness, accompanied by an increase in the on/off current ratio (fig.5d). In addition, g_m_ values show a progressive increase from 2.7±0.3 mS for l3 to 3.5±0.4 mS for l2 to 4±0.2 mS for l1 (fig.5e) underscoring the significant influence of channel length on the amplification abilities of OECTs. Although less pronounced to changes in g_m_ which nearly doubled for a 3-time decrease in length, the response time improved from 1±0.1 ms for l3 to 0.8±0.02 ms for l1 (fig.5f and fig.5g) leading to a subsequent rise in the cut-off frequency from 160 Hz to 200 Hz (fig.5h). This evaluation of the channel length highlights its impact on OECT performance and offers some insights for consideration of OECT designs for all intended applications which will be discussed later.

### Label-Free Electrochemical Sensing of NT-proBNP with Functionalized Inkjet OECTs

In this section, we use the W500, l2, and d2 variation, for its high amplification (gm = 12.5 mS) and moderate speed, transitioning to an all-planar architecture by inkjet patterning a 6.25 mm2 gold gate contact near the channel resulting in a small form factor, requiring only a few microliters (enough to cover the gate and the channel) to operate, as shown in figure 6a. This electrode, functionalized with monoclonal antibodies (fig.6b), enables NT-proBNP detection with a gate capacitance 41.83 µF ± 34.75 nF, and a gate-to-channel capacitance ratio of approximately 5.22, ensuring effective gating ^43^. FTIR confirmed successful antibody functionalization, displaying the two distinct vibrational bands in 1700-1850 and 1580-1650 cm-1, representing the amide I and II bands (fig.6c). These bands are the two signatures of protein backbones, the earlier is caused by C=O stretch and the latter is caused by the N-H and C-N stretching vibration. The transmittance of these bands intensifies after each immobilization step due to higher proteins density on the surface. Device I-V and transfer characteristics post functionalization show no hindrance in device performance, but a shift in threshold voltage to from 0.4 V to 0.5 V (fig.6d) and a consequent shift in operating point from 0.26 V to 0.31 V (fig.6e), pre and post functionalization respectively. This shift is attributed to the increased negative charges on the gate surface. However, this shift is not significant enough to push the device to an operating region outside the water window of PEDOT:PSS (−0.9 V to 0.6 V) or Gold (−0.6 V to 0.8 V) ^44^. At pH 8.5, the NT-proBNP proteins gain a negative charge (isoelectric point = 7.23), and upon binding to the gate through the immobilized antibodies, the effective gate-source voltage decreases, increasing drain current (figure 6c). As shown in figure 6d, the device exhibits linear sensitivity from 10 to 400 pg/ml, with a limit of detection of 68 pg/ml and a resolution of 0.038% ΔIDS/pg/ml, saturating above 500 pg/ml. This range is clinically relevant, as NT-proBNP levels ≥125 pg/ml indicate early-stage B heart failure with a 2-fold increased risk of cardiovascular mortality^45,46^. The red shaded area in figure 6f is the device’s response to buffer only control sample aliquots. Our findings position inkjet-printed OECTs as scalable, cost-effective platforms for field-deployable NT-proBNP sensors, with potential for multiplexed cardiac biomarker panels in decentralized clinical settings. Material costs for a single OECT are $0.15, with functionalization adding $1.38, yielding a total cost of $1.53 per unit.

**Figure 6.**
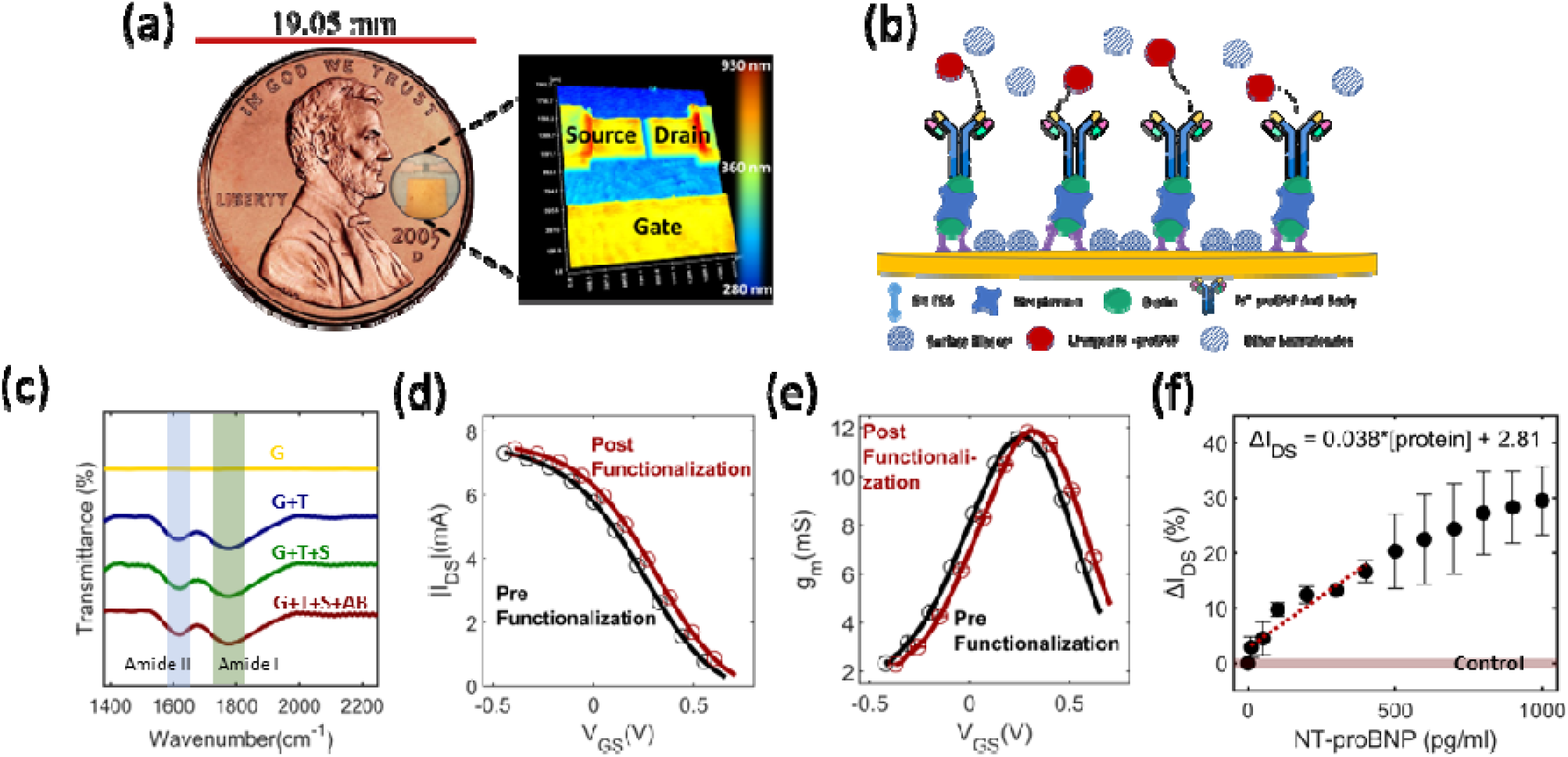
Inkjet OECTs to detect heart failure biomarkers. (a) small OECT footprint allows the use of small sample size to operate. (b) sketch of the functionalization steps to immobilize antibodies on the gate surface. (c) FTIR spectra highlighting the increase of amide I and II magnitudes with every functionalization step. (d) transfer characteristic pre and post functionalization. (e) g_m_ curve pre and post functionalzation. (f) percent of drain current shift function of NT-proBNP concentration from 0 to 1000 pg/ml

### Flexible Organic Transistors for Dynamic Electrocorticography in Epilepsy Models

In this section, we evaluate the performance of the W100, d1, and l1 device for biopotential recording, focusing on its speed and amplification for high signal-to-noise ratio (SNR) measurements. We first validated its performance in a benchtop experiment by recording a simulated hippocampal neuron population spike injected at the gate. As shown in Supplementary Figure 4, the device accurately captures low-amplitude signals (∼100 µV) with a 25 ms temporal profile, achieving an SNR of 31.9 dB. These results underscore the OECT’s low noise and high amplification, making it ideal for low amplitude biopotential applications. We then demonstrate the device’s capability to monitor seizure progression in vivo using a rat model. Our previous ECoG study showed that OECTs outperform low-impedance passive electrodes, highlighting the advantages of active thin-film organic transistor technology ^41^. Here, we focus on the ability of these inkjet OECTs to track epileptic episodes. Seizures were induced by kainic acid injection into the amygdala, and a flexible OECT was placed on the somatosensory cortex. Recording began 23 minutes post-injection, with continuous data collected until minute 50 (Supplementary Figure 5). Initially, recordings resemble those under ketamine anesthesia, displaying low-frequency, low-amplitude oscillations. Spontaneous high-frequency, elevated-amplitude spikes (∼100 µV above baseline, fig.7) emerge around 27–27.5 minutes, increasing in regularity by minutes 32–33. These spikes grow in frequency until a major epileptic episode occurs at approximately minute 44, characterized by high-amplitude (∼350 µV) polyspike bursts (maximum of 70 Hz) and slow waves (1Hz) lasting ∼30 seconds. Post-episode, spontaneous lower-amplitude spikes persist for ∼5 minutes, followed by a second kainic acid-induced seizure signature; continuous high-amplitude (∼400 µV), high-frequency (maximum of 80 Hz) spikes lasting ∼15 seconds. Towards the end of our recordings and prior to sacrificing the animal, we see a return to baseline activity with spontaneous high frequency spikes. As demonstrated by our animal model, these findings confirm that flexible inkjet-printed OECTs are highly effective for in vivo monitoring of seizure progression, reaching an SNR up to 48 db, on par with previously reported OECTs ^47,48^.

**Figure 7.**
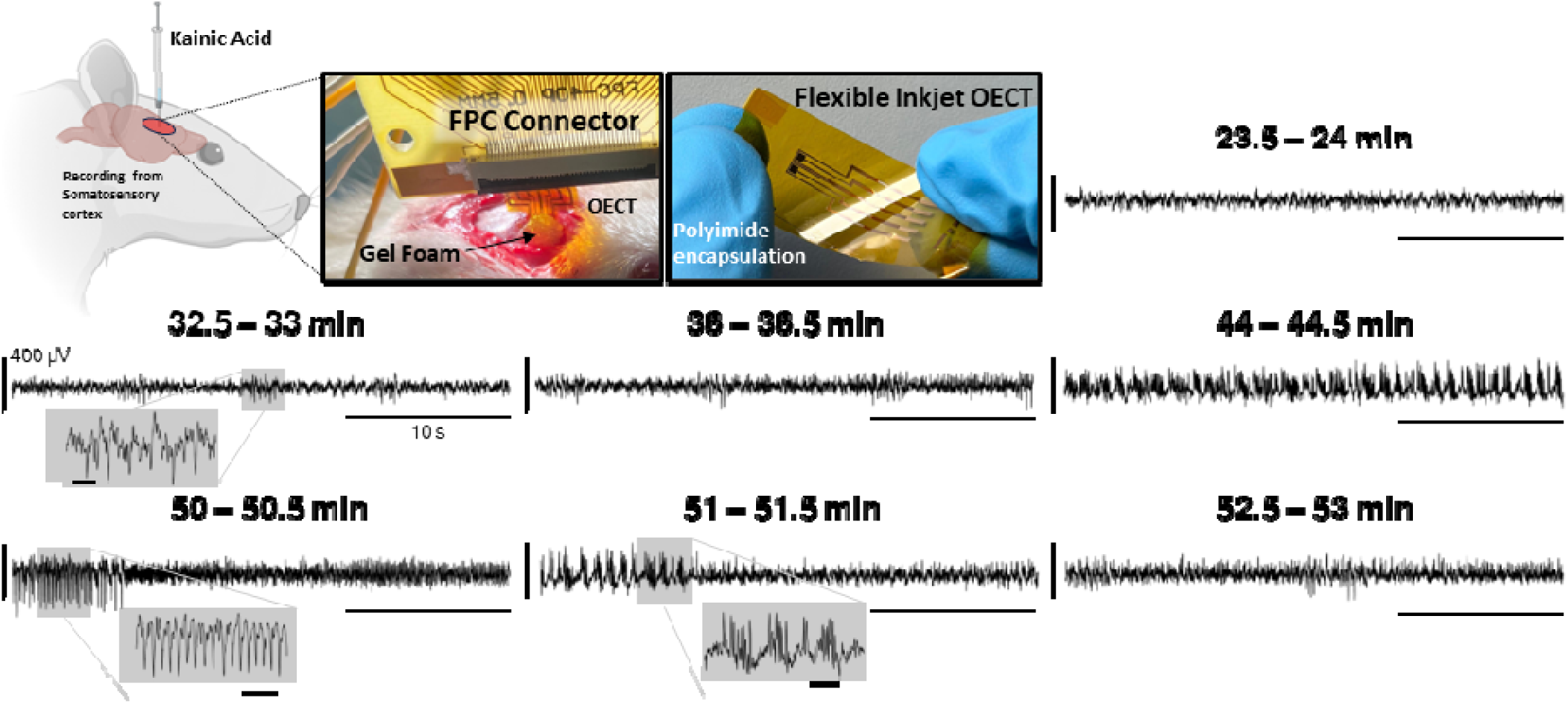
In vivo recording of kainic acid induced seizure in rats using flexible inkjet printed OECTs. Minute 23.5-24 is a representative recording under anesthesia pre seizure. Minutes 32.5 and 36 represent recordings of spontaneous high frequency spikes. Minutes 44 and 51 are representative of the burst and slow wave pattern. Minute 50 is representative of the high frequency, high amplitude continuous spikes. Scale bars in the grayed-out insets are 20ms.

## Discussion

Through a methodical examination of the Inkjet microfabrication limits and how device geometrical parameters affect OECT performance, we aimed to highlight the potential in realizing high performing OECTs with high stability and reproducibility. The latter is essential to meet the requirements of bioelectronic interfaces for sensing and electrophysiological applications. Inkjet printing brings versatility and precision to fabricating OECTs, enabling tailored designs for bioelectronic interfaces while eliminating complex cleanroom processes. Its precise material deposition allows for optimal morphological tuning, including footprint, thickness, and uniformity, while accommodating various substrate materials (rigid and flex). Our technique leverages oxygen plasma treatment to modify the surface wettability, resulting in parameters that surpass previously reported dimensions for printed OECTs. As demonstrated in Table 1 and compared to other printed OECT work, we achieved the smallest channel lengths and thickness while maintaining one of the smallest channel widths (fig. S6a). Furthermore, our performance metrics indicate that our work boasts transconductance values among the highest reported, as well as the lowest response times by a significant margin (fig. S6b). Additionally, our technique resulted in low error bars across fabrication trials, demonstrating consistent results both in fabrication and electrical performance. Because the scalability, yield, and reproducibility of Inkjet OECTs has been a continuous problem^49^, our improvements in this study not only enhance their performance and applicability but also paves the way for their integration into more complex and miniaturized electronic systems.

**Table 1.**
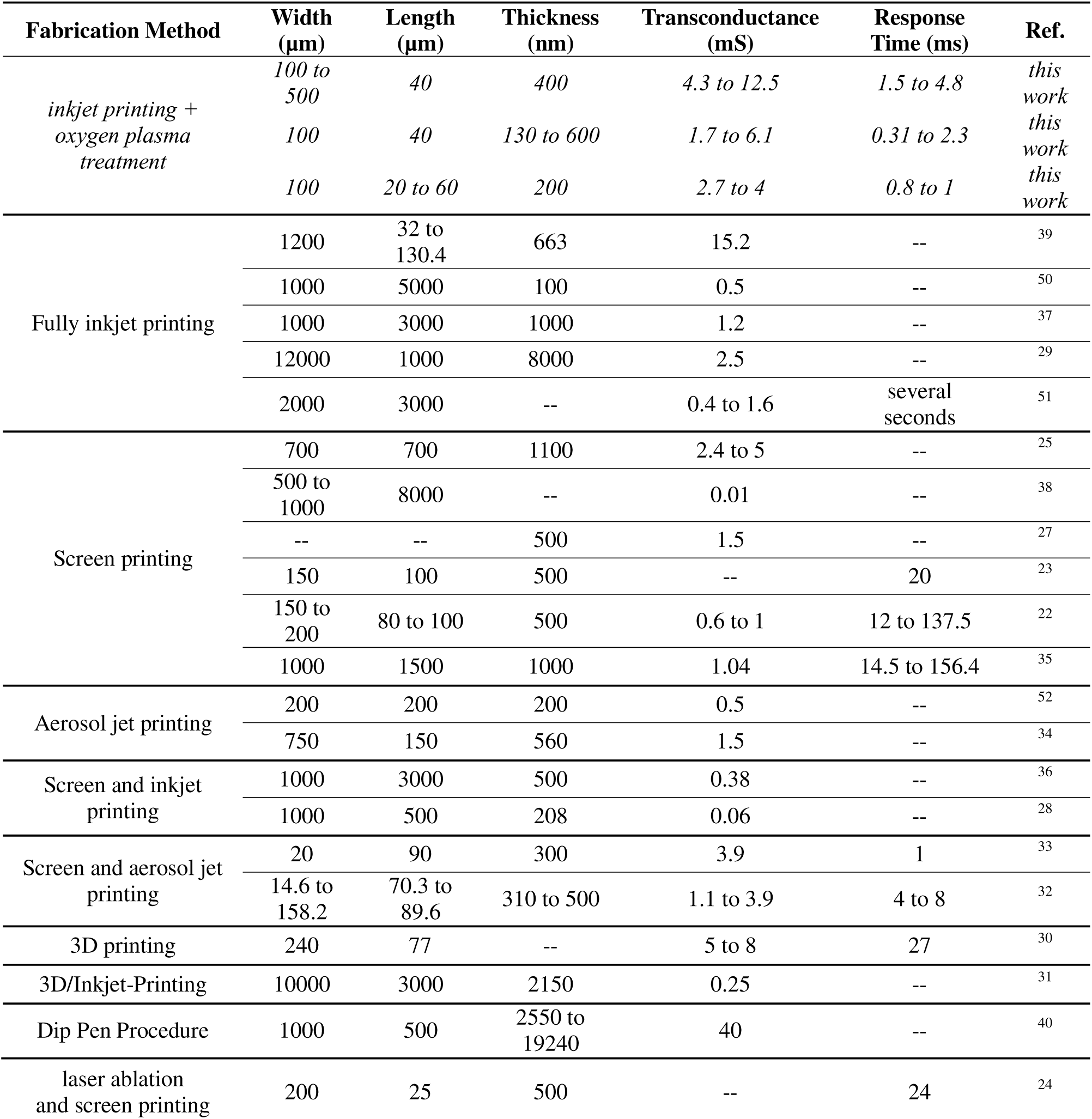
Comparison of printed OECT geometries and key performance metrics.

### For biomolecule detection applications

Adequate width scaling of OECTs presents a viable strategy to enhance the amplification capabilities at the expense of response time; a parameter less critical in biosensors engineered for biomolecule detection, where the target molecules have very low concentration, requiring high amplification to detect their presence. For instance, a response time of 4.8 ms for W500 devices is deemed sufficient considering it has a g_m_ of 12.5mS. Widening the channel was found to significantly enhance the on/off ratio, indicating improved switching capability. This observed increase amplifies the SNR, thereby increasing sensitivity and reducing the limit of detection^18^. Another important aspect to achieve reliable detection and signal amplification is being able to control V_GS, gm max_ which can be effectively managed by adjusting the channel width. These adjustments offer significant advantages for redox chemical transducers for example by employing the gate at their redox potential^53^. Importantly, the stability of inkjet-printed OECTs in solution for molecular sensing is crucial for stable and reliable operation across transfer characteristic scans, as the detection of the target is commonly quantified by the degree of shift from a baseline “empty” scan. Our W500 device achieved notably high stability through the initial 50 scans after which g_m_ increased, coupled with a lateral shift in maximum g_m_ and a drop in the drain current. This can be attributed to PEDOT:PSS’s parasitic electrochemical reactions and microstructure disruption in solution over time. The minimal error across all 100 scans (equivalent to 40 µA) coincided with V_GS, gm max_ and amounts to only 0.86% of the drain current (I_DS, gm max_ = 4.65 mA). These results underscore precision, with deviations below the typical shifts seen in target-sensitive OECTs for DNA ^54^ or proteins ^55^ sensing, affirming the reliability of V_GS, gm max_ to be the point at which target monitoring should be evaluated across time. Furthermore, inkjet micropatterning permits a seamless integration of a planar gate electrode serving as the site on which the target can bind through chemical surface functionalization, making OECTs sensitive and specific to the desired biomolecule. We use such technique in this study to immobilize monoclonal antibodies specific to NT-proBNP, a circulating heart failure biomarker, resulting in a biosensor with a detection range of 10 to 400 pg/ml with a resolution of 0.038% / pg/ml. These metrics are especially important in a clinical setting, as this range is of interest to assess the risk of heart failure.

### For biopotential recording

OECTs are distinguished by their superior amplification capabilities, yet they encounter significant performance declines at high frequency signals, due to the inability of ions to penetrate the full bulk of the channel at high speeds, concentrating the channel doping at the uppermost surface, and essentially transforming the device’s behavior to a conventional field effect transistor, thereby reducing g_m_ ^56,57^. This phenomenon poses a challenge for the integration of OECTs into electrophysiological applications, requiring speeds up to 1 kHz. Nonetheless, decreasing the channel’s size (whether in width, thickness, or length) has been proven to mitigate the adverse effects of high frequencies on the gain of the device; and with inkjet printing, it is possible to easily tune these parameters. Also, understanding brain micromotion–induced strain is critical for designing durable, low-stress neural interfaces; computational models have demonstrated how strain magnitude and spatial distribution around intracortical implants can guide substrate flexibility and mechanical compliance. While inkjet printing offers very precise control in a 2D plane, its ability to modulate the thickness of the deposited ink is inherently bound to a discreet number of layers. Despite these limitations, the channel thickness parameter can still be adjusted to an extent by carefully modulating the DS, a measure of drop-to-drop distance as depicted by the schematic in figure 4a. This control is however, significantly impacted by the contact angle of the ink with the substrate^58^. A high contact angle necessitates a smaller DS to achieve uniform material deposition, while too low an angle requires a larger DS. Therefore theoretically, the minimum achievable channel thickness is dictated by the ink-substrate contact angle. We overcome this constraint by introducing oxygen plasma treatment to graft the surface with hydroxyl groups, thus enhancing its wettability, and consequently, decrease the contact angle. Two approaches were adopted to achieve thinner depositions without compromising the print’s size. The first involved preserving the DS and extending the plasma treatment time, thereby reducing the volume of ink required to achieve the desired print size and resulting in much thinner channels (fig.4b). Alternatively, the second method involved brief oxygen plasma treatment of the glass substrate combined with high DS to prevent print gap formations (fig.4c). In fact, our realized plasma treatment-complemented device is sufficiently quick to decode EEG, ECoG, LFPs, and ECG. However, accurately representing faster electrophysiological signals like action potentials and EMG signals are beyond the capabilities of the current OECT configuration^15^. Further device performance tuning is possible by varying the length of the channel. This strategy emerges as an effective approach as modifying channel length - unlike thickness and width - can uniformly enhance the OECT performance on all aspects without detriments on amplification or response time. Therefore, decreasing length to the limit of the microfabrication technique adopted is desired to maximize performance on all fronts irrespective of the application. In inkjet techniques for fabricating OECTs, the lowest achievable length matches the set DS, in our case 20 µm (or 1 drop-difference), which is generally on par with high-performing non-inkjet OECTs^59^. Modifying wettability prior to the metal deposition could further allow smaller channel length by achieving controlled spreading akin to what’s been proposed for thickness, thereby pushing the limits of what can be achieved with inkjet. As a demonstration of its recording capability, we utilize inkjet printing to fabricate devices on a flexible polyimide substrate, allowing conformality to the brain. Using a seizure animal model, we were successfully able to detect seizure progression, from low amplitude anesthesia-specific patterns to higher amplitude and frequency spike seizure behavior.

All in all, this work shows the potential for inkjet printing in the fabrication of high-performance OECTs with high precision and stability, enabling the fabrication of devices with various g_m_ values, scaling up with (fig. S7a), in addition to the control of response time down to a optimization data, our results place this technique as a viable option for the fabrication of OECTs, matching their photolithographed counterparts in both figures of merit ^11,15^.

The limitations of our fabrication lie in the need to optimize printable fluids to eject from the nozzles, and the dependency of drop morphology, spreading, and solvent evaporation on room temperature and humidity. There are significant opportunities in the microfabrication of inkjet-printed OECTs, including the ability to accommodate a wide range of substrate types. However, a notable threat is its resolution; although sufficient for most applications, it cannot achieve resolutions below 15 micrometers. A comprehensive SWOT analysis depicted in supplementary figure 7c highlights the pivotal considerations for inkjet-OECTs, underpinning their potential impact and future opportunities.

## Conclusion

OECTs are traditionally fabricated using photolithography multi-step fabrication techniques. In this study, we explore the use of cold fabrication inkjet printing in the development of OECTs. Comparatively to clean room fabrication, inkjet reduces material waste, is environmentally friendly, is scalable, and is compatible with unconventional substrates including biocompatible flexible polymers with low melting points^60, 61^. Also, understanding brain micromotion–induced strain is critical for designing durable, low-stress neural interfaces; computational models have demonstrated how strain magnitude and spatial distribution around intracortical implants can guide substrate flexibility and mechanical compliance^62^. These findings align with earlier efforts on microscale resonant biosensors for neuronal adhesion, which similarly emphasize the importance of mechanical and surface properties in neural interface design^63^. Oxygen plasma surface treatment prior to deposition allowed the realization of performance metrics otherwise unachievable and expanded the horizon of possibilities for printed-OECTs, offering transconductance ranges of up to 15 mS and response times down to 0.31 ms. This work also showcases the flexibility of inkjet printing in adjusting OECT designs seamlessly to suit specific application needs. We demonstrate the latter by using one variation of these OECTs to detect heart failure biomarkers with a resolution of 0.038% / pg/ml in the 10 to 400 pg/ml range, and another variation was used to monitor seizure progression in a rat animal model, capturing transitions from anesthetic baseline to epileptic activity. To conclude, this paper serves as a practical guide for fabricating OECTs using inkjet printing, offering a cost-effective and efficient alternative to cleanroom methods. These advancements pave the way for broader adoption of Inkjet OECTs in biomedical diagnostics and neuromonitoring.

## Data availability statement

All data will be made available upon reasonable request.

## Author contributions

Fadi Khoury: Conceptualization, Methodology, Software, Validation, Formal analysis, Investigation, Data Curation, Writing - Original Draft, Writing - Review & Editing, Visualization, Project administration.

Zeina Habli: Data Curation, Methodology, Validation, Investigation, Writing – Review & Editing.

Jad Daorah: Investigation, Data Curation. Yuchen Xu: Writing – Review & Editing.

Makram Obeid: Supervision, Writing – Review & Editing.

Samir Alam: Conceptualization, Writing - Review & Editing, Supervision. Gert Cauwenberghs: Supervision, Writing – Review & Editing.

Massoud Khraiche: Conceptualization, Resources, Writing - Review & Editing, Supervision, Project administration, Funding acquisition.

## Supporting Information

Supporting information is available from the Wiley Online Library.

