## Supplementary for "Geometry-Optimized Inkjet-Printed Organic Electrochemical Transistors with High Transconductance and Sub-Millisecond Response for Biosensing and Neural Recording"

**Supplementary Material**

Advancing the Inkjet Microfabrication of Organic Electrochemical Transistors for Biomarker Detection and Neural Recordings

Fadi Khoury^1^, Zeina Habli^1^, Jad Daorah^1^, Yuchen Xu^2^, Makram Obeid^3^, Samir Alam^4^, Gert Cauwenberghs^2^, and Massoud Khraiche^1,*^

^1.^Neural Engineering and NanoBiosensors Group, Biomedical Engineering Program, Maroun Semaan Faculty of Engineering and Architecture, American University of Beirut, Beirut 1107 2020, Lebanon.

^2.^Department of Bioengineering, Jacobs School of Engineering, University of California San Diego, La Jolla, CA 92093, USA.

^3.^Stark Neurosciences Research Institute, Department of Neurology, Indiana University School of Medicine, Indianapolis, Indiana 46202, USA.

^4.^Department of Internal Medicine, American University of Beirut, Beirut 1107 2020, Lebanon.

### **Notes for supplementary figure 1:**

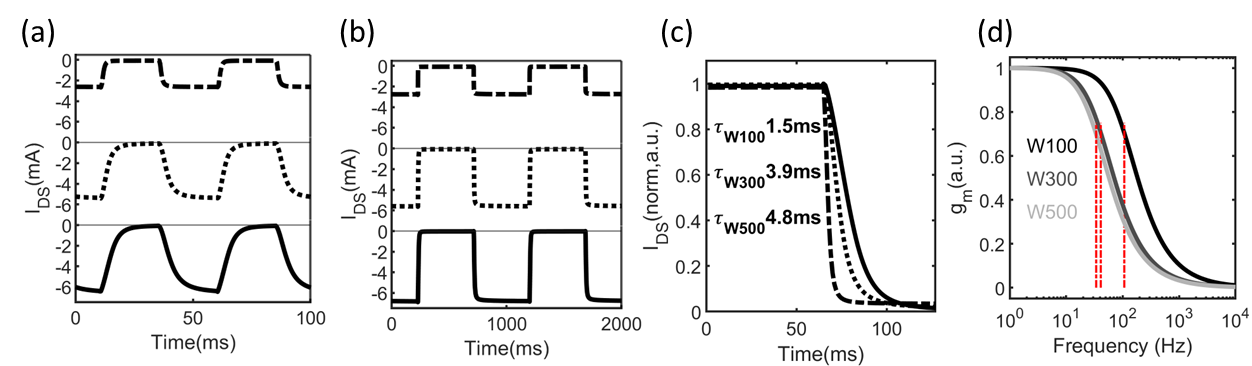

Supplementary Figure 1 (a,b) current response of W100 (top), W300 (middle), and W500 (bottom) to a (a) 20Hz and (b) 1Hz square wave applied at the gate. (c) zoomed in view of the current transition from high to low and the corresponding response time. (d) amplitude drop of gm function of frequency for W100 (black), W300 (grey), and W500 (light grey)

Increasing the width of the active channel decreases the speed of OECTs, represented by the response time. However, to correctly extract this value, it is essential to supply the square wave at an adequate frequency so that the devices can correctly reconstruct it. Figure S2a is an example of the three different OECT configurations responses to a frequency that is too quick to be fully reconstructed (20Hz). Figure S2b is the response of the same devices at a lower square wave frequency (1Hz), from which response time can be accurately extracted as shown in Fig.S2c, and consequently cut-off frequency can be deduced as depicted in Fig.S2d.

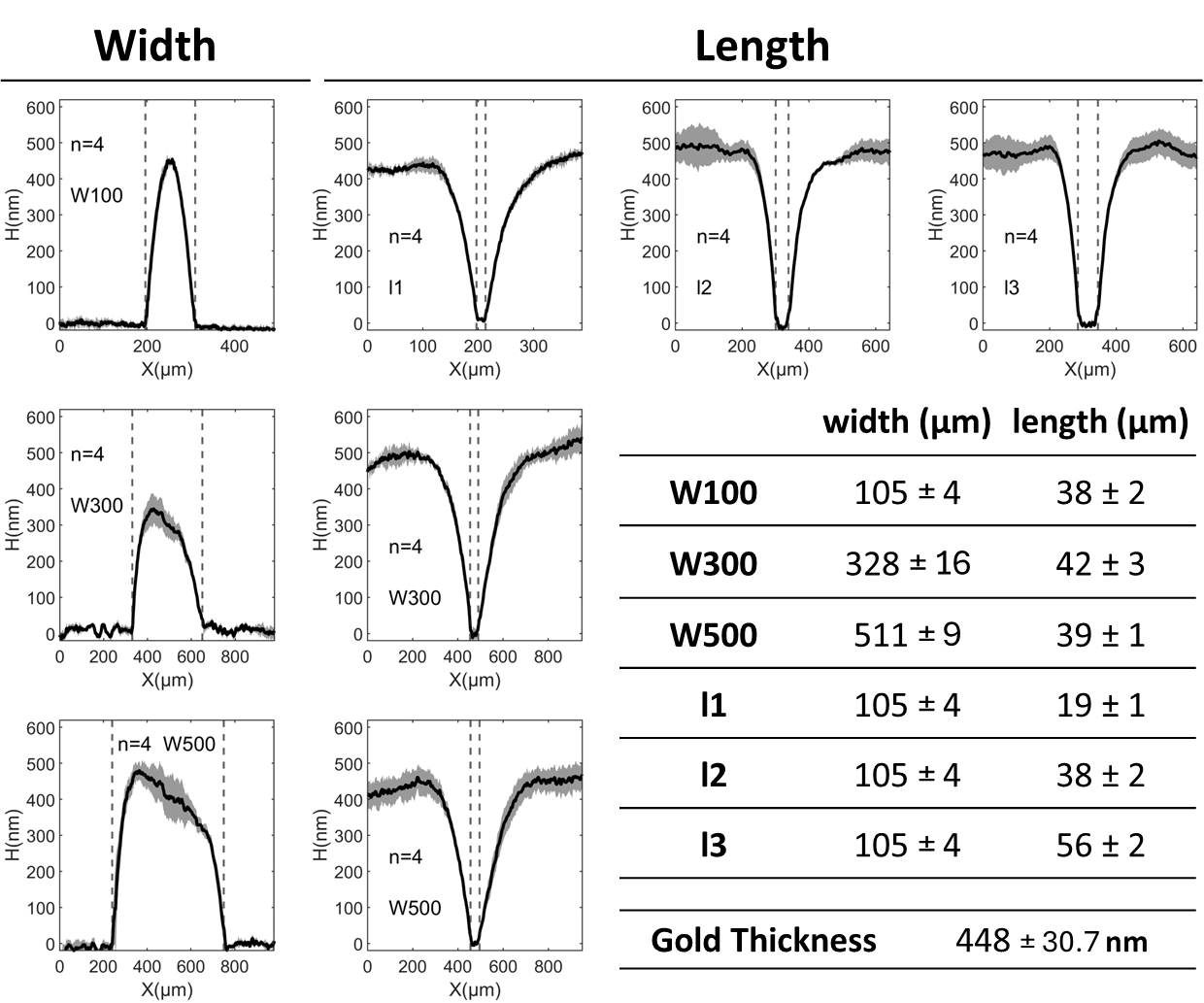

Supplementary Figure 2 DHM analysis of the fabricated widths and lengths using inkjet printing

### **Notes for supplementary figure 3:**

The use of the complementary oxygen plasma treatment technique facilitated the production of OECTs with very thin channel thicknesses. Fig.S3a corresponds to averaged IV curves for produced devices revealing very low error bars across fabrication iterations, proving the method’s reliability and reproducibility. The advantage of this additional technique is that it allowed for very low response times down to 0.31ms, as seen in Fig.S3b, unachievable otherwise.

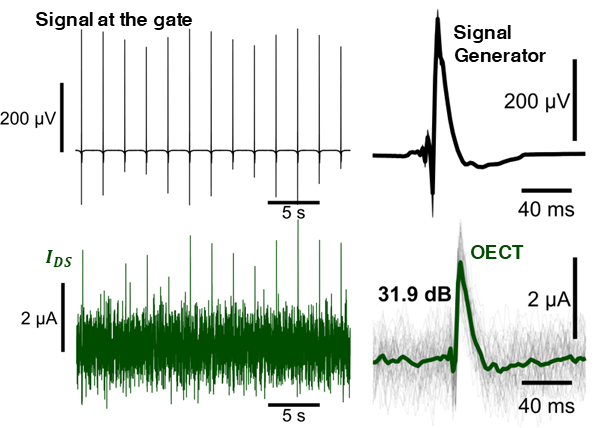

Supplementary Figure 4 Benchtop recording (OECT in green) of artificial neuronal spikes injected at the gate (black)

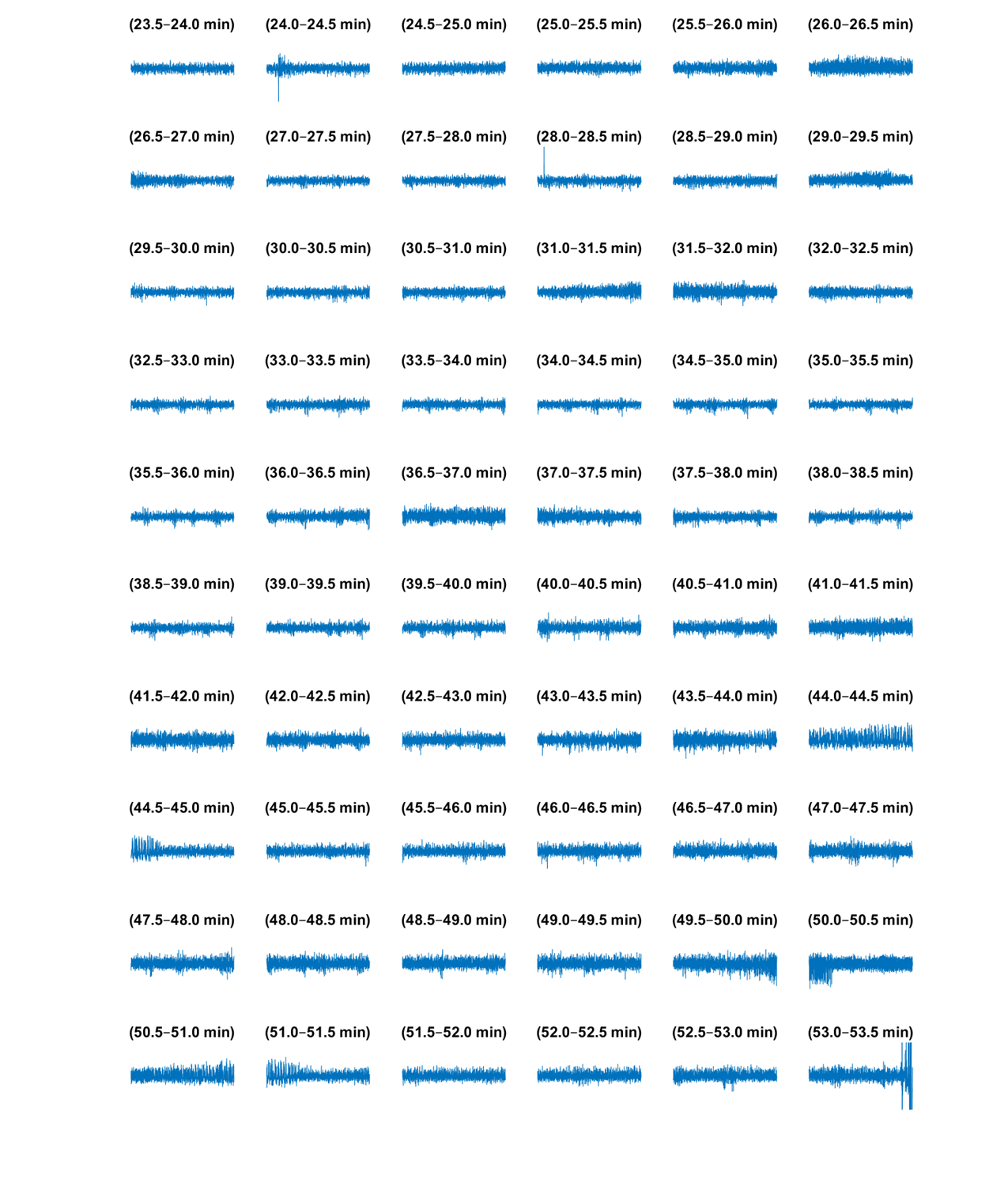

Supplementary Figure 5 Full OECT ECoG recordings during seizure.

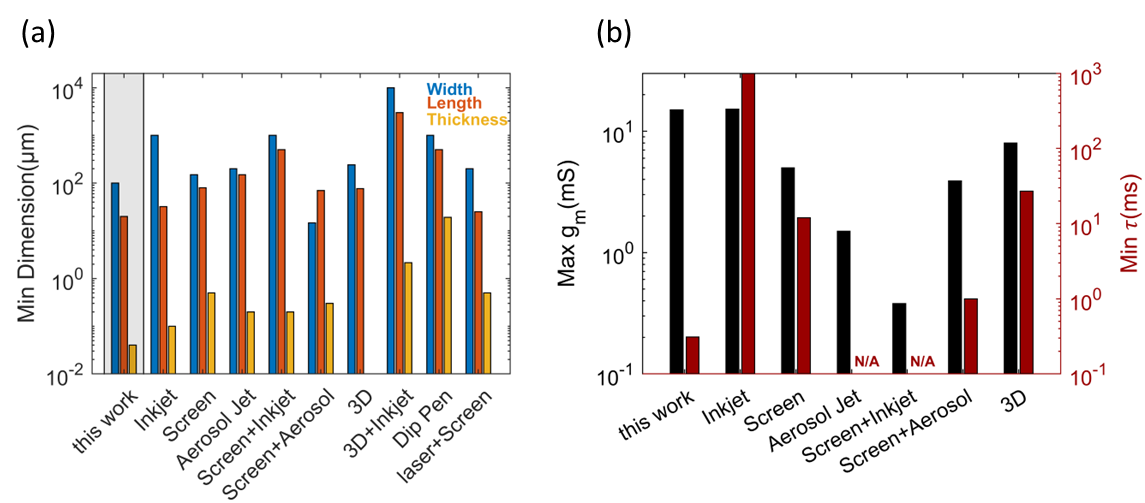

Supplementary Figure 6 (a) minimal OECT widths, lengths, and thicknesses achieved by various printing methods. (b) maximal and minimal OECT transconductances and response times, respecively, achived by various printing methods.

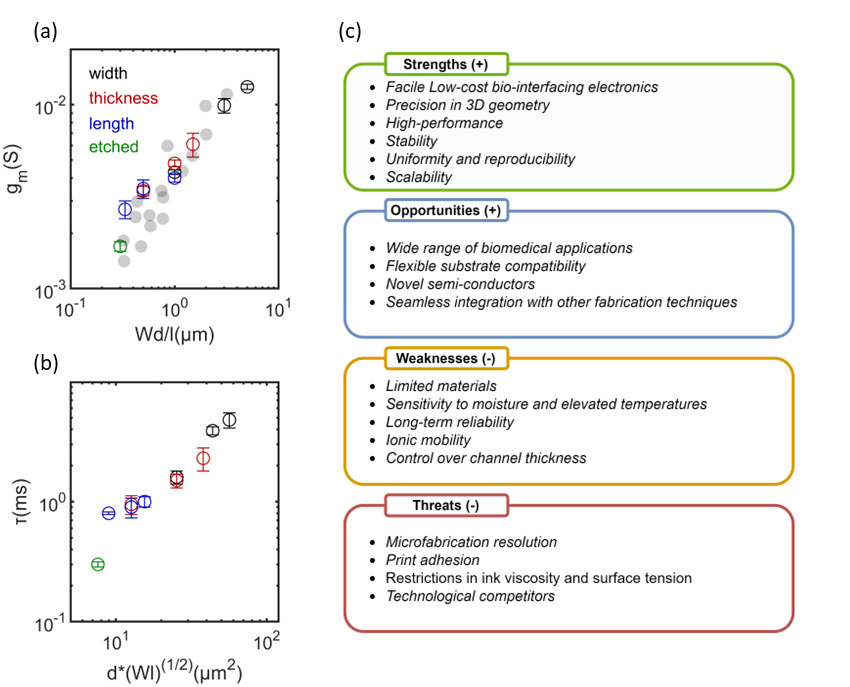

Supplementary Figure 7 Control of (a) gm and (b) τ through tuning of width (black), thickness (red), length (blue), and etch-complemented (green). Grey data points in (a) are adopted from [12] with permission under a Creative Commons Attribution NonCommercial License 4.0 (CC BY-NC). 10.1126/sciadv.1400251. (c) Strength, Weakness, Opportunities, and Threats (SWOT) of inkjet-printed OECTs.
